# Does the neuronal noise in cortex help generalization?

**DOI:** 10.1101/676999

**Authors:** Brian Hu, Jiaqi Shang, Ramakrishnan Iyer, Josh Siegle, Stefan Mihalas

**Affiliations:** Allen Institute for Brain Science

## Abstract

One remarkable feature of neuronal activity in the mammalian cortex is the high level of variability in response to repeated stimuli. First, we used an open dataset, the Allen Brain Observatory, to quantify the distribution of responses to repeated presentations of natural movies. We find that even for their preferred moment in the movie clip, neurons have high variability which cannot be well captured by Gaussian or Poisson distributions. A large fraction of responses are better fit by log-normal or Gaussian mixture models with two components. These distributions are similar to activity distributions during training of deep neural networks using dropout. This poses the interesting hypothesis: is the role of cortical noise to help in generalization during learning?

Second, to ensure the robustness of our results we analyzed electrophysiological recordings in the same areas of mouse visual cortex, again using repeated natural movie presentations and found similar response distributions. To make sure that the trial-by-trial variations we observe are not due exclusively to the result of changes in state, we constructed a population coupling model, where each neuron’s activity is coupled to a low-dimension version of the activity of all other simultaneously recorded neurons. The population coupling model can capture global, brain-wide activity fluctuations that are state-dependent. The residuals from this model also show non-Gaussian noise distributions.

Third, we ask a more specific question: is the noise in the cortex more likely to move the representation of the stimulus in-class versus out-of-class? To address this question, we analyzed the responses of neurons across trials from multiple sections of different movie clips. We observe that the noise in the cortex better aligns to in-class variations. We argue that a useful noise for learning generalizations is to move from representations of different exemplars in-class, similar to cortical noise.

## 1 Introduction

One of the hallmarks of neuronal codes is the high level of trial-to-trial variability [1, 2]. It should be noted that this variability is predominantly in the central nervous system, as peripheral fibers can be surprisingly precise [3]. The trial-to-trial variability of cortical activity has been studied using multiple stimuli [4], as well as its relation to attention [5] and other behavioral variables [6]. Previous theories on the possible role of noise center on its potential usefulness in inference [7].

The goal of this study is not to be exhaustive in the characterization of cortical noise. Even an exhaustive review of the literature would be hard, but it is aided by several review papers [7, 8]. Rather, the goal of the present study is to present one surprising observation which came out of analysing the response distributions in a large-scale survey of neural activity in the awake mouse visual cortex conducted across multiple visual areas and cell types [9], and to perform hypotheses-driven experiments and analyses which follow-up on this observation.

There are three sections to this study: in the first part, we analyze response distributions in a large-scale survey of neural activity in the awake mouse visual cortex conducted across multiple visual areas and cell types [9]. The surprising observation is that very few cells are fit by a Gaussian model, even for their preferred stimulus. Most are best fit by either log-normal or two-component Gaussian mixture models, most often with one of the mixtures near zero. A Gaussian mixture model with two components, one of which is at zero, is one observed characteristic of units within networks trained with “dropout”, a model which has been shown to prevent model overfitting and reduce feature co-adaptation [10]. Dropout assumes neurons can be modeled as independent and identically distributed Bernoulli random variables, and randomly “drops out” neurons within a network with some probability *p*.

The optical signal from two-photon calcium fluorescense is not a direct measurement of spiking activity. The observation of the response distribution at the level of calcium dynamics in the soma being non-Gaussian can be interpreted in different ways depending on how the distributions are transformed through the spiking nonlinearity and captured in electrophysiological recordings. To address this question, in the second part of the study, we performed electrophysiological recordings in three mice (see Section 4) using repeated presentations of natural movies. Performing a similar analysis as the optical physiology, we obtain qualitatively similar results.

An additional question is whether the observed log-normal and two-component Gaussian mixture distributions are induced by brain-wide state changes. However, defining the state based purely on the observation of behavior is difficult. We cannot address this question directly, but we instead use a more indirect means: a recent study [6] observed that the behavioral state of the animal is reflected well in a dimensionally-reduced representation of neural activity recorded across multiple areas of the central nervous system. If the non-Gaussian distributions are exclusively the result of state fluctuations, they should be captured by the coupling of the activity of individual cells with this lower-dimensional population activity. Towards this end, we used a dimensionality reduction method (see section 4) and constructed models linearly coupled with this low-dimensional state. We rerun our analyses on the residuals obtained from this model. We again observe the same log-normal and Gaussian mixture models using the residual activity, leading us to conclude that brain state cannot exclusively explain the variability in neural activity.

**These observations lead us to formulate the central hypothesis of the paper: that networks of cortical neurons use noisy representations with the goal of constructing meaningful categories for the elements present in images, and that this noise allows for building of general representations from a small number of exemplars.**

Using this assumption of the underlying computation, if only one category is needed, the ideal noise takes the representation of one exemplar in one category and maps it to all other exemplars with probabilities equal to occurrences in the organisms’ environment. The relevant features for categorization might also vary with the task at hand. However, if enough categories have similar axes of variations along relevant features, the ideal noise would move the representation along axes which are often conserved in a category. We do not know exactly which axes these are, but if we pick etologically relevant categories, our hypothesis is that the noise of an exemplar in a category should be better aligned to variations in that category rather than other categories. However for the mouse visual system, etologically relevant categories are hard to define. For our analyses, we used movie clips randomly selected from a set of natural movies. This analysis is the focus of the third part of the study.

## 2 Results

### 2.1 Noise distribution in Allen Brain Observatory data

We analyzed the variability in neural activity across visual areas, layers, and transgenic mouse lines using data from the Allen Brain Observatory [9]. Specifically, we analyzed neural responses to one of the presented natural movie stimuli (a 30-second clip from the opening scene of the classic movie “Touch of Evil”). Our hypothesis is that structure in the response distributions provides insight into how neural representations change on a trial-to-trial basis can help us build hypothesis about their role. For each cell, we identified its preferred “stimulus” within the movie clip and quantified trial-to-trial variability by fitting different distributions to the neural responses across trials (*N* = 30) for the cell’s preferred “stimulus” (see Methods for details). Figure 1C,D shows the variability in neural responses for an example cell in our dataset.

**Figure 1:**
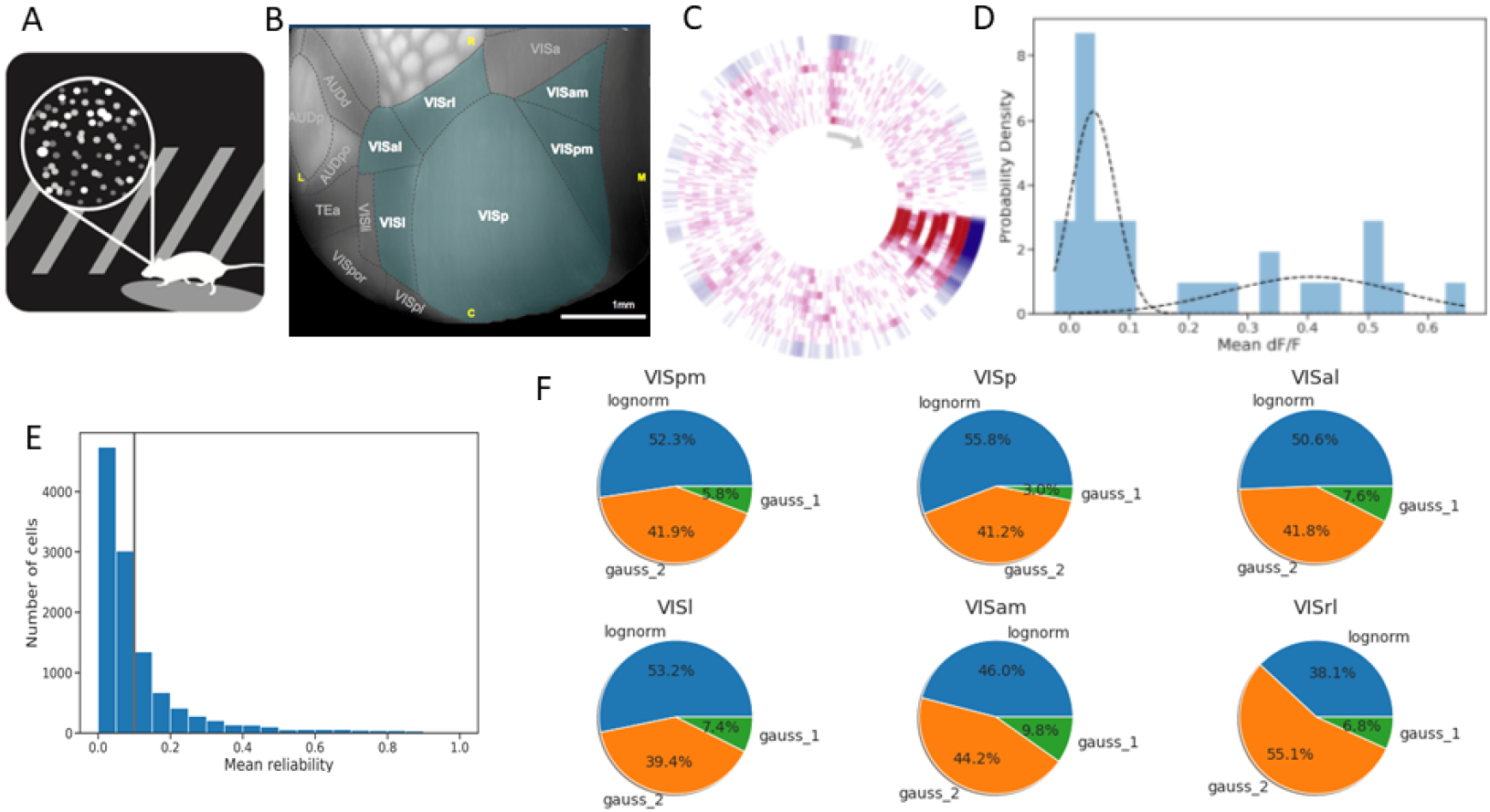
Neural variability in the Allen Brain Observatory [9]. (A) These analyses are based on two-photon optical recordings. (B) Anatomy of the visual areas in which recordings were performed. (C) Track plot visualization of neural responses to the natural movie stimulus in the Allen Brain Observatory for an example cell. Frames of the movie are shown clockwise starting with the gray arrow, with the ten repeats within a session shown in red extending radially. The mean response across trials is shown by the outer blue ring. For many frames of the movie, the cell does not respond on every trial, even though the stimulus shown is exactly the same. (D) The distribution of responses for an example cell across all trials (three sessions with ten repeats each). The two components of the Gaussian mixture model are shown overlaid in the dotted black lines. (E) Distribution of response reliabilities for all cells in the dataset. We set a minimum reliability threshold of 0.1 (gray vertical line), and only characterized the response distributions of cells above this threshold. (F) Percent of cells in each of the recorded visual areas, broken down by the best-fit response distribution. Distributions include log-normal and one- and two-component Gaussian mixtures.

One measure of response reliability is the mean trial-to-trial correlation of neural activity within a session, which is bounded between 0 (low reliability) and 1 (high reliability) [9]. Over the entire population of cells, we find very low mean response reliabilities across sessions, with a mean reliability of 0.11 (Figure 1E). To remove extremely unreliable cells from our analysis, we set a reliability threshold of 0.1. Our subsequent analyses are performed using this set of cells (*N* = 1775), which has a mean reliability of 0.37. We find that the vast majority of cells are better fit by either log-normal distributions or Gaussian mixture models with two components. To determine the best-fitting response distribution, we performed model selection using the Akaike information criterion (AIC), but using other information theoretic measures yielded similar results. An additional analysis based on bootstrap parametric cross-fitting comparing one component and two-component Gaussian mixture models confirmed the robustness of our model selection procedure (Supplementary Info). Figure 1F summarizes the response distribution fits across cells in our dataset. We found consistent results across layers and mouse transgenic lines which labeled excitatory cells (Supplementary Figure 3). Untangling cell-type specific differences in neural variability (including in different inhibitory cells) and their contributions to learning is an important area of future research.

We also performed a similar analysis using the units within a convolutional neural network. For this analysis, we trained a simple network with dropout on the CIFAR-10 image dataset [11]. Normally, dropout is only used during training, and turned off during evaluation. Here, we continued to use dropout during evaluation, which introduces variability in the responses of each unit. Using dropout in this way has been proposed as a form of Bayesian approximation [12]. Critically, we find that the bimodal distribution in neural responses across trials can also be captured by a convolutional neural network trained with dropout (Supplementary Figure 1).

### 2.2 Noise distribution and state dependence in electrophysiological recordings

To test if the noise distributions observed in optical physiology also exist in spiking activity, we also analyzed the variability in neural activity across multiple visual areas using spiking data obtained with high-density Neuropixel probes from 3 separate experiments. Specifically, we analyzed spiking responses in neurons (936 total units across all visual areas and mice) to 98 repeats of natural movie stimuli consisting of several distinct movie clips (see Section 4).

We used the same methods as for optical physiology to define the preferred frame in the movie for each cell, and found qualitatively similar results: across areas and layers, most cells are better fit by log-normal or Gaussian mixture models with two components. This result persists even when including Poisson and negative binomial distributions as alternative hypotheses to be tested (Poisson and negative binomial are not good distributions to test the continuous *dF/F* signals from optical physiology).

To control for potential state changes we performed an additional analysis and estimated the noise distribution for each neuron as follows. We isolated each neuron and estimated its coupling to the instantaneous network state using the activities of all the remaining simultaneously recorded neurons (population coupling). We obtained a reduced (100-dimensional) representation of the instantaneous network state by hierarchically clustering the activities of the remaining neurons into 100 clusters (see Methods for details). We used the average activity of neurons in these 100 clusters and fit a linear model that uses these 100 inputs to predict the activity of the single neuron of interest. We computed residuals for each neuron using the difference between the observed responses and the predicted model responses.

We quantified trial-to-trial variability by fitting different distributions to the neural response residuals across trials (*N* = 98) for the cell’s preferred “stimulus” (see Methods for details). As with the imaging data, we find that the vast majority of cells are still better fit by either Gaussian mixture models with two components or log-normal distributions. This was true even after incorporating Poisson and negative binomial as candidate distributions (Figure 2).

**Figure 2:**
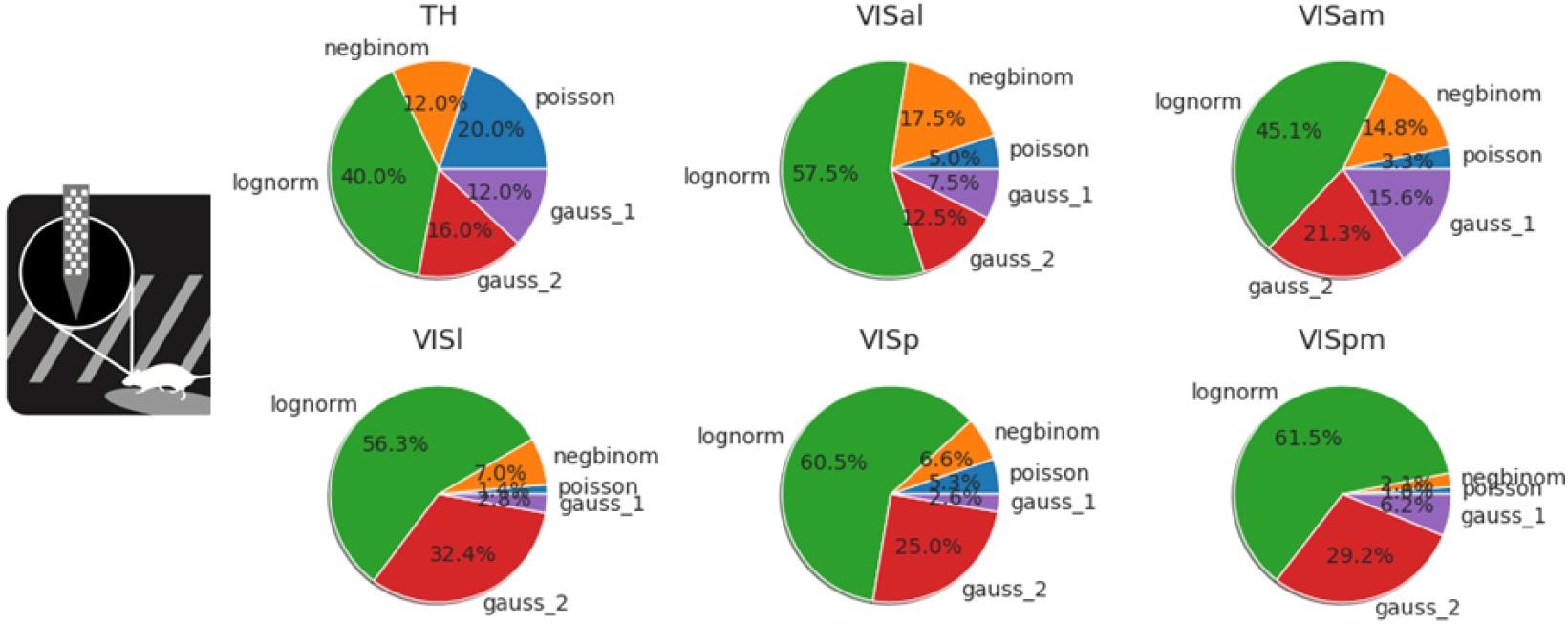
Quantification of neural variability to the preferred stimulus in cells across visual areas from electrophysiology data. To control for possible effects of state, we fit a model coupling the neuronal responses to a dimensionally reduced population activity of the other neurons. We fit the noise distribution on the residuals of the model. We also observe a large fraction of cells across visual cortical areas being best fit by log-normal or a two component Gaussian Mixture model. The only area where a non-trivial fraction of neurons are best fit by Poisson is subcortical, visual thalamus.

### 2.3 Trial-by-trial variability mimics in-class exemplar changes

We quantify how similar the noise in representation of one exemplar is to: 1) variations in representations of exemplars in the same or other clips, 2) typical representations of the same or other clips, and 3) typical representations of exemplars in the same or other clips. The presence of non-Gaussian noise makes the use of traditional methods to tackle this question difficult to interpret. For example, dropout-like noise produces a projection in the neuronal activity subspace which cannot be captured by a Gaussian distribution. As a result, we proceed with a set of very simple measures based on the neuronal activity subspace.

We define the activity for neuron *i* of an exemplar *j* in clip *k* in trial *n* as the spike count during the exemplar time window *a*_*i,j,k,n*_ (see Section 4). The signal for neuron *i* of exemplar *j* in clip *k* is the average over trials of the activity *s*_*i,j,k*_ = 〈*a*_*i,j,k,n*_〉_*n*_. The noise for neuron *i* of an exemplar *j* in clip *k* in trial *n* is the activity minus the signal *n*_*i,j,k,n*_ = *a*_*i,j,k,n*_ − *s*_*i,j,k*_.

We define several subspace measures based on the neuronal activity.

- The exemplar coding subspace for an exemplar and clip is defined as the set of neurons for which the signal is bigger than the average *E*_*j,k*_ = (*s*_*i,j,k*_ > 〈*s*_*i,j,k*_〉_*j,k*_).
- The clip coding subspace is defined as the set of neurons for which the signal is larger than the mean for more than half of the exemplars in the clip, *C*_*k*_ = (*mean*_*j*_(*s*_*i,j,k*_ > 〈*s*_*i,j,k*_〉_*j,k*_) *>*= 0.5).
- The clip variance subspace is defined as the set of neurons for which the variance of signal for exemplars in the clip is larger than for exemplars across clips, *V*_*k*_ = (*std*_*j*_(*s*_*i,j,k*_) > *std*_*j,k*_(*s*_*i,j,k*_)).
- The noise subspace for an exemplar, clip, and trial is defined as the set of neurons for which the absolute value of the noise is larger than its standard deviation. *N*_*j,k,n*_ = (*abs*(*n*_*i,j,k,n*_) *> std*_*j,k,n*_(*n*_*i,j,k,n*_)). We also explored the possibility of defining the absolute value of the noise to be larger than a constant times its standard deviation. Since the results are robust to the exact value of the constant, we chose its value to be 1.

First we used a custom similarity metric, projection similarity *P* (*S*1*, S*2), which measures how many of the axes of a vector *S*2 are in the vector *S*1. This is inspired by the intuition that noise variations of an exemplar in clip *k*_1_ should be a subset of the clip *k*_1_ coding subspace but not clip *k*_2_ as depicted in Figure 3A. We computed this measure for each mouse and each visual area with at least 20 reliable neurons. The pooled distribution is observed in Figure 3B. The differences are highly statistically significant (p<0.001). What is more impressive is that this result is in the right direction and is statistically significant (p<0.05) in each individual area analyzed (number of areas in all mice = 8).

**Figure 3:**
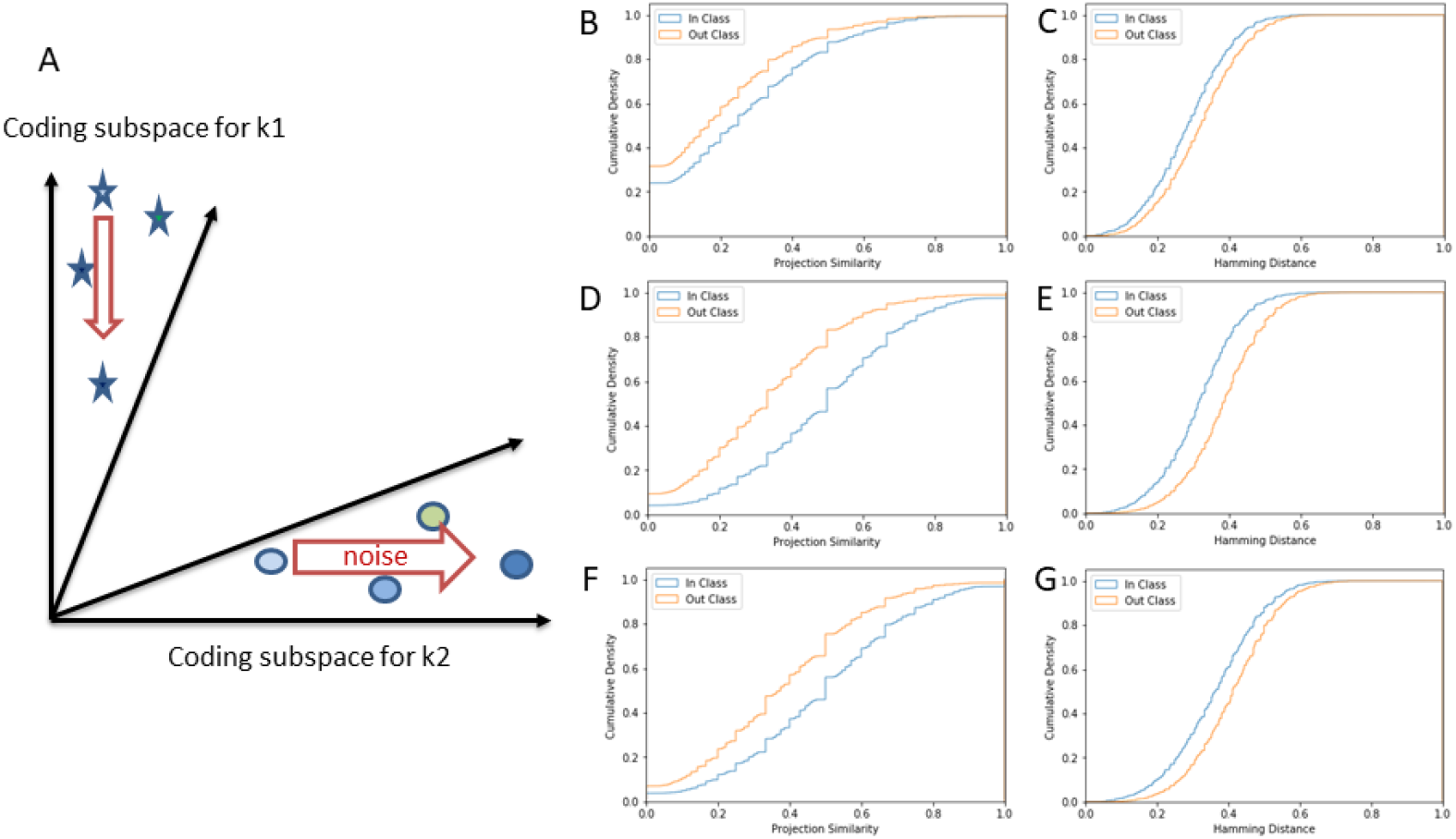
Quantification of direction of neural variability in the full population of cells across visual areas. (A) The main thesis of the paper: noise in the cortex for an exemplar is aligned with the coding subspace for the category it belongs to (clip coding subspace) and to the variations in the coding subspace between exemplars in the same category (clip variance subspace). (B-G) Cumulative distributions for similarity metrics using data aggregated across 3 mouse recordings. Projection similarity (B) and Hamming distance (C) between clip variance subspace *V*_*k*_ and noise subspace *N*_*j,k,n*_. Projection similarity (D) and Hamming distance (E) between clip coding subspace *C*_*k*_ and noise subspace *N*_*j,k,n*_. Projection similarity (F) and Hamming distance (G) between exemplar coding subspace *E*_*k*_ and noise subspace *N*_*j,k,n*_. We observe closer alignment of the noise subspace to each of these subspaces within the same clip versus across different clips.

To make sure that the results are not entirely dependent on the similarity metric used, we repeated the analysis using a more standard Hamming distance (Figure 3 C). The Hamming distance between two binary vectors measures the numbers of dimensions which do not agree. The distance between the noise subspace for exemplars in clip *k*_1_ are smaller to the clip *k*_1_ variance subspace than to the clip *k*_2_ variance subspace. The differences are again highly statistically significant (p<0.001). Just as for the projection similarity, this result is in the right direction and is statistically significant (p<0.05) in each individual area analyzed.

While we can not fully address the algorithms or mechanisms of how the cortical network can generate noise for each exemplar which better aligns to the same clip subspace variance, the observation that it also aligns with the same clip coding subspace (Figure 3 D, E) and the exemplar coding subspace (Figure 3 F, G), allows us to speculate on a potential mechanism (see Discussion below).

## 3 Discussion

In the first and second part of the paper we observed complex, non-Gaussian distributions of noise in neurons’ coding for their preferred stimulus. The role of noise for generalization in artificial neuronal networks has been explored (see [10], and the wealth of citations in that study). While we drew inspiration from neurons with dropout-like noise to generate the hypothesis about generalization, it does not mean that other noise distributions do not contribute to generalization. We do not know enough about variation of neuronal activity across exemplars of multiple etologically relevant categories to draw a firm conclusion on the optimality of different distributions. It might be possible that different types of categorization would preferentially use different noise distributions.

The non-Gaussian distributions observed inspired the use of a subspace analysis at the neuronal level. As dropout-like noise generates a projection in neuronal space, eliminating some neurons altogether, it is the most natural place to focus the analysis of alignment. While the number of mice used in our study is low, the results are surprisingly strong from a statistical significance point of view. The alignment of trial-to-trial noise of an exemplar in a clip is statistically significantly better to the same clip exemplar-by-exemplar variation than for other clips, for every visual area in every individual mouse we tested.

This study is not aimed at addressing algorithmically or mechanistically how the noise which is aligned with in-class variations arises. Though we can speculate that as the network does align the in-class variations with the class code, a somewhat simpler mechanism in which trial-to-trial variability of an exemplar aligns with typical representation of that exemplar, can lead to the computation observed.

It would be very interested if this phenomenology can be replicated in primates, within which the etologically relevant axes of generalization are better known. We believe that research into new forms of biological noise can also be used to train neural networks with better generalization capabilities. Extensions of dropout which address the fact that neurons are often highly correlated with each other have been proposed [13, 14], but we believe that inspiration from biological noise and the computations implemented could be a very fruitful avenue for improving generalization capabilities from low numbers of examples.

## 4 Methods

### 4.1 Allen Brain Observatory Data

We used the “natural movie one” stimulus from the Allen Brain Observatory. The movie clip was presented across three different imaging sessions, with ten repeats within each session, for a total of 30 trials. A total of *N* = 11731 cells had measurable reliabilities across all three imaging sessions. We set a minimum reliability threshold of 0.1 across each of the three sessions to eliminate extremely unreliable cells from our analysis. This eliminated many of the cells, and resulted in a subset of *N* = 1775 cells. For our subsequent analyses, we first broke the movie clip into 1-second epochs (each consisting of 30 frames), and computed the mean *dF/F* response over each epoch. To identify the “preferred” stimulus for each cell, we computed the epoch which elicited the max mean *dF/F* response over all epochs and all trials. This provides us with a distribution of responses across each cell’s “preferred” stimulus within the movie.

### 4.2 Noise distribution fitting

For each cell, we quantified the distribution of neural responses across trials by fitting different distributions using the *scikit-learn* package. The distributions we fit included one- and two-component Gaussian mixtures, log-normal, Poisson, and negative binomial. We only fit Gaussian mixture and log-normal distributions to the Allen Brain Observatory data, which is based on continuous *dF/F* responses. The Poisson and negative binomial distributions require discrete counts, which could only be applied to the electrophysiological recording data. We used the Aikaike information criterion (AIC) to select between model fits, although other information theoretic measures yielded qualitatively similar results. For our dropout noise analyses, we used a bootstrap parametric cross-fitting test with *N* = 10, 000 samples to determine the significance of the two-component Gaussian mixture model fits. This test effectively compares how likely the difference in log-likelihoods between the one- and two-component models can be achieved purely by chance, given the null hypothesis that the data is generated from a single component distribution. For cells which were deemed better fit by the two-component Gaussian mixture model, we performed an additional test to determine whether their response distributions were dropout-like. For each of these cells, we calculated a z-score on the component with the lower mean, and those cells with z-scores less than two (meaning their means are not significantly different than zero) were counted as cells with dropout-like response distributions. The number of cells that pass these two tests divided by the total number of cells gives the fraction of cells which have dropout-like response distributions. We also performed additional analyses, separating cells by visual area, layer, and transgenic mouse line.

### 4.3 Neural network training with dropout

To test whether our neural response distributions match those from a neural network trained with dropout, we used a simple network architecture. We trained a neural network with four convolutional layers and two fully-connected layers on the CIFAR-10 image dataset. During training and evaluation, we used a dropout percentage of 0.5. As we were not concerned with state-of-the-art image classification accuracy, we trained our model for only ten epochs using stochastic gradient descent with a learning rate of 0.01 and minibatch size of 64. To produce variable response distributions, we presented the network with the 118 natural images used in the Allen Brain Observatory. We computed the “preferred” stimulus for each unit within the network by finding the stimulus that evoked the max mean activation across 50 repeats of each image.

### 4.4 Electrophysiological recordings

#### Animal preparation

All experimental procedures were approved by the Allen Institute for Brain Science Institutional Animal Care and Use Committee. Five weeks prior to the experiment, mice were anesthetized with isoflurane, and a metal headframe with a 10-mm circular opening was attached to the skull with Metabond. In the same procedure, a 5-mm-diameter craniotomy and durotomy was drilled over left visual cortex and sealed with a circular glass coverslip. Following a 2-week recovery period, a visual area map was obtained through intrinsic signal imaging [15]. Mice with well-defined visual area maps were gradually acclimated to the experimental rig over the course of 12 habituation sessions. On the day of the experiment, the mouse was placed under light isoflurane anesthesia for 40 min to remove the glass window, which was replaced with a 0.5 mm thick plastic window with laser-cut holes (Ponoko, Inc., Oakland, CA). The space beneath the window was filled with agarose to stabilize the brain and provide a conductive path to the silver ground wire attached to the headpost. Any exposed agarose was covered with 10,000 cSt silicone oil, to prevent drying. Following a 1-2 hour recovery period, the mouse was head-fixed on the experimental rig. Up to six Neuropixels probes coated in CM-DiI were independently lowered through the holes in the plastic window and into visual cortex at a rate of 200 m/min using a piezo-driven microstage (New Scale Technologies, Victor, NY). When the probes reached their final depths of 2,500–3,500 m, each probe extended through visual cortex into hippocampus and thalamus.

#### Data acquisition

In vivo recordings were performed in awake, head-fixed mice allowed to run freely on a rotating disk. During the recordings, the mice passively viewed a battery of visual stimuli, including local drifting gratings (for receptive field mapping), full-field flashes, and 100 repeats of an 80 s natural movie stimulus. All spike data were acquired with Neuropixels probes [16] with a 30-kHz sampling rate and recorded with the Open Ephys GUI [17]. A 300-Hz analog high-pass filter was present in the Neuropixels probe, and a digital 300-Hz high-pass filter (3rd-order Butterworth) was applied offline prior to spike sorting.

#### Data preprocessing

Spike times and waveforms were automatically extracted from the raw data using Kilosort2 [6]. Kilosort2 is a spike-sorting algorithm developed for electrophysiological data recorded by hundreds of channels simultaneously. It implements an integrated template matching framework for detecting and clustering spikes, rather than clustering based on spike features, which is commonly used by other spike-sorting techniques. After filtering out units with “noise” waveforms using a random forest classifier trained on manually annotated data, all remaining units were packaged into Neurodata Without Borders format [18] for further analysis.

### 4.5 Population coupling model

To characterize the coupling of each neuron to the population activity of the other simultaneously recorded neurons, we isolated this neuron and first clustered the single-trial activity of the remaining neurons in to 100 clusters using hierarchical clustering. This was done using the agglomerative clustering method provided by the scikit-learn package with average linkage and the pairwise Pearson correlation coefficient of single-trial activities. We calculated the average activity of neurons within each cluster and used the 100 signals thus generated as a predictor for the single-trial activity of a single neuron in a linear model. Linear model fitting was performed using GLM tools provided by the *statsmodels* package with the Gaussian family with an identity link function. We split the single-trial activities into two equal halves, using the first half as a training set and the second half as the test set. The difference between the model response and the experimentally observed neuronal response was used to calculate the residual activity for each neuron. The residual activities were then binned into the same shape as the single-trial neuronal responses (number of neurons × number of movie frames × number of trials) and were fitted using the methods outlined in the section above.

### 4.6 In-class and out-of-class movie analysis

We analyzed the activity during 11 natural movie clips ranging from 4 to 9 seconds each for a total of 81s were selected at random from large database of natural movies and were repeated 98 times. As in the optical physiology, we only analyzed neurons and trials with a reliability above 0.1. We did this analysis inside a visual area for each mouse separately if at least 20 reliable neurons were collected in each (9 areas pass this threshold). The activity during the first 1 sec following a clip transition was not used during this analysis (although tests including it show similar results). The analysis starts by randomly choosing 10 non-overlapping exemplars from each clip (the results are robust to this choice), which are 200 ms long sections (200 ms was chosen for this presentation as it is longer than typical autocorrelation of single neuron activity when injected with a current in the soma, but results are robust to changes in this parameter from 33ms to 1s when dropping the number of exemplars to ensure non-overlap). The signal for each exemplar is the spike count of each neuron during the exemplar movie section.

