## Supplemental Material for "Does the neuronal noise in cortex help generalization?"

Supplementary Material

#### Additional Analysis of Allen Brain Observatory Data

As an alternative to the AIC method for testing best fit distributions, we supplement the results with a bootstrap test. For each cell, we identified its preferred "stimulus" within the movie clip and quantified trial-to-trial variability by fitting Gaussian mixture models to the distribution of responses across trials ($N = 30$) for the cell's preferred "stimulus".

For each cell, we quantified the distribution of neural responses across trials by fitting Gaussian mixture models with either one or two components to the distributions using the *scikit-learn* package. We used a bootstrap parametric cross-fitting test with $N = 10,000$ samples to determine the significance of the two component model fits. This test effectively compares how likely the difference in log-likelihoods between the one and two component models can be achieved purely by chance, given the null hypothesis that the data is generated from a single component distribution.

We find that 88.7% of the cells (10403/11731 cells) are better fit by Gaussian mixture models with two components compared to one component, indicating a two component Gaussian Mixture Model better fir the distribution in their responses compared to one component ($p < 0.05$, bootstrap parametric cross-fitting test). Model selection based on information theoretic measures such as AIC resulted in a similar proportion of cells with two component response distributions (92.0%, 10794/11731 cells). Using only a subset of the cells with reliabilities above a minimum reliability threshold (see below for details) resulted in a lower, but still substantial proportion of cells (80.6%, 1430/1775 cells). Figure 1 shows the variability in neural responses for an example cell in our dataset.

We also performed a similar analysis using the units within a convolutional neural network. For this analysis, we trained a simple network with dropout on the CIFAR-10 image dataset [1]. As this model was trained with static images rather than movies, we instead performed our analyses using the set of 118 natural images in the Allen Brain Observatory. We passed each natural image through the model 50 times, and recorded the distribution of activations for each unit within the model to its "preferred" stimulus. Normally, dropout is only used during training, and turned off during evaluation. Here, we continued

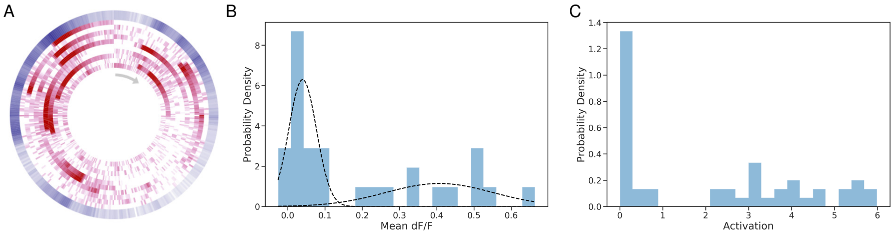


Figure 1: Neural variability for an example cell from the Allen Brain Observatory and an example unit within a neural network trained on the CIFAR-10 image dataset. (A) Track plot visualization of neural responses to the natural movie stimulus in the Allen Brain Observatory for an example cell (session A, cell specimen id 517447794). Frames of the movie are shown clockwise starting with the gray arrow, with the ten repeats within a session shown in red extending radially. The mean response across trials is shown by the outer blue ring. For many frames of the movie, the cell does not respond on every trial, even though the stimulus shown is exactly the same. (B) The distribution of responses for the example cell in (A) across all trials (three sessions with ten repeats each). The two components of the Gaussian mixture model are shown overlaid in the dotted black lines. (C) The distribution of activations for an example unit within a neural network (second convolutional layer, fourth feature map) across all trials (50 repeat presentations of natural images).

to use dropout during evaluation, which introduces variability in the responses of each unit. Using dropout in this way has been proposed as a form of Bayesian approximation [2]. Critically, we find that the bimodal distribution in neural responses across trials can also be captured by a convolutional neural network trained with dropout (compare Fig 1b and Fig 1c).

We also quantified neural variability at the population level. One measure of response reliability is the mean trial-to-trial correlation of neural activity within a session, which is bounded between 0 (low reliability) and 1 (high reliability) [3]. Over the entire population of cells, we find very low mean response reliabilities across sessions (Figure 2a), with a mean reliability of 0.11. To remove extremely unreliable cells from our analysis, we set a reliability threshold of 0.1. Our subsequent analyses are performed using this set of cells ($N = 1775$), which has a mean reliability of 0.37. To identify cells with dropout-like response distributions, we first found the cells that were better fit by Gaussian mixture models with two components, and for each of these cells, we computed a z-score on the component with the lower mean to test whether it was significantly different from zero. For cells which were deemed better fit by the two-component Gaussian mixture model, we performed an additional test to determine whether their response distributions were dropout-like. For each of these cells, we calculated a z-score on the component with the lower mean, and those cells with z-scores less than two (meaning their means are not significantly different than zero) were counted as cells with dropout-like response distributions. The number of cells that pass these two tests divided by the total number of cells gives the fraction of cells which have dropout-like response distributions. We also performed additional analyses, separating cells by visual area, layer, and transgenic mouse line. Based on this measure, we find that the fraction of cells with dropout-like response distributions is high and relatively constant across visual areas, with a mean fraction of 0.74 (Figure 2b). We also find only modest differences in the fraction of cells with dropout-like distributions across layers and mouse transgenic lines, with the lowest fraction of cells being in the superficial layers (Figure 2c). Finally, we also studied the noise distributions across Cre-lines in our dataset (Figure 3). We found consistent results across all Cre-lines, which selectively labeled excitatory cells in different cortical areas and layers.

### References


[1] Alex Krizhevsky. Learning multiple layers of features from tiny images. 2009.

[2] Yarin Gal and Zoubin Ghahramani. Dropout as a bayesian approximation: Representing model uncertainty in deep learning. In *international conference on machine learning*, pages 1050–1059, 2016.

[3] Saskia E J de Vries, Jerome Lecoq, Michael A Buice, Peter A Groblewski, Gabriel K Ocker, Michael Oliver, David Feng, Nicholas Cain, Peter Ledochowitsch, Daniel Millman, Kate Roll, Marina Garrett, Tom Keenan, Leonard Kuan, Stefan Mihalas, Shawn Olsen, Carol Thompson, Wayne Wakeman, Jack Waters, Derric Williams, Chris Barber, Nathan Berbesque, Brandon Blanchard, Nicholas Bowles, Shiella Caldejon, Linzy Casal, Andrew Cho, Sissy Cross, Chinh Dang, Tim Dolbeare, Melise Edwards, John Galbraith, Nathalie Gaudreault, Fiona Griffin, Perry Hargrave, Robert Howard, Lawrence Huang, Sean Jewell, Nika Keller, Ulf Knoblich, Josh Larkin, Rachael Larsen, Chris Lau, Eric Lee, Felix Lee, Arielle Leon, Lu Li, Fuhui Long, Jennifer Luviano, Kyla Mace, Thuyanh Nguyen, Jed Perkins, Miranda Robertson, Sam Seid, Eric Shea-Brown, Jianghong Shi, Nathan Sjoquist, Cliff Slaughterbeck, David Sullivan, Ryan Valenza, Casey White, Ali Williford, Daniela Witten, Jun Zhuang, Hongkui Zeng, Colin Farrell, Lydia Ng, Amy Bernard, John W Phillips, R Clay Reid, and Christof Koch. A large-scale, standardized physiological survey reveals higher order coding throughout the mouse visual cortex. *bioRxiv*, 2018.

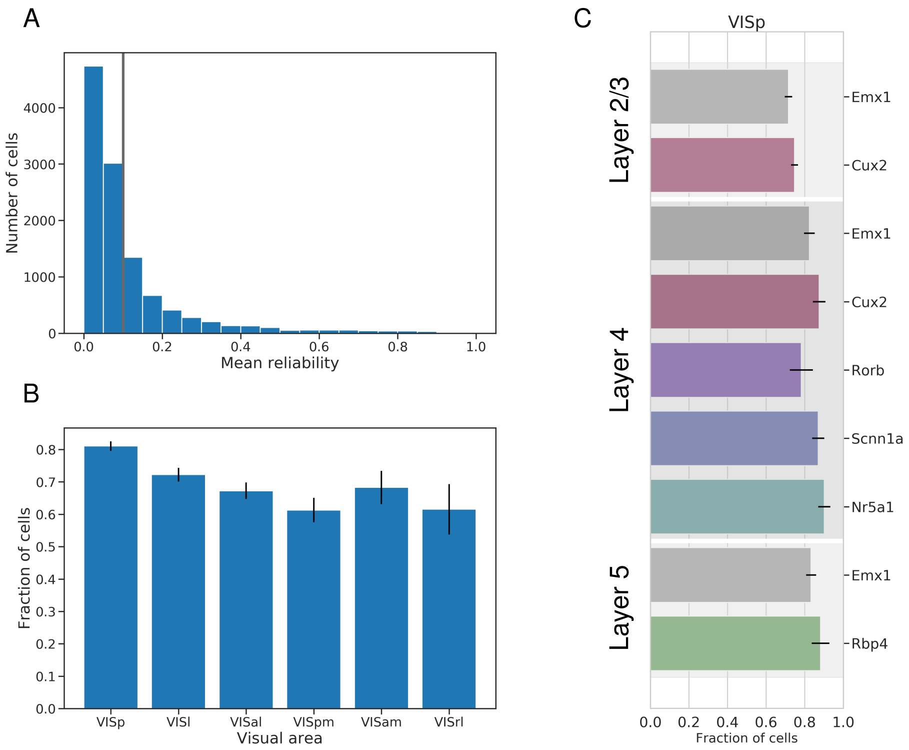


Figure 2: Quantification of neural variability in the full population of cells across visual areas, layers, and transgenic mouse lines. (A) The distribution of mean reliabilities (average trial-to-trial correlations) across the three imaging sessions for all cells included in our analysis. The vertical gray line shows the reliability threshold criterion of 0.1 which we used. (B) The fraction of cells with response distributions consistent with dropout across visual areas (mean = 0.74). Error bars show standard deviations. (C) The fraction of cells with distributions consistent with dropout across layers and transgenic mouse lines in primary visual cortex (VISp). Error bars show standard deviations.

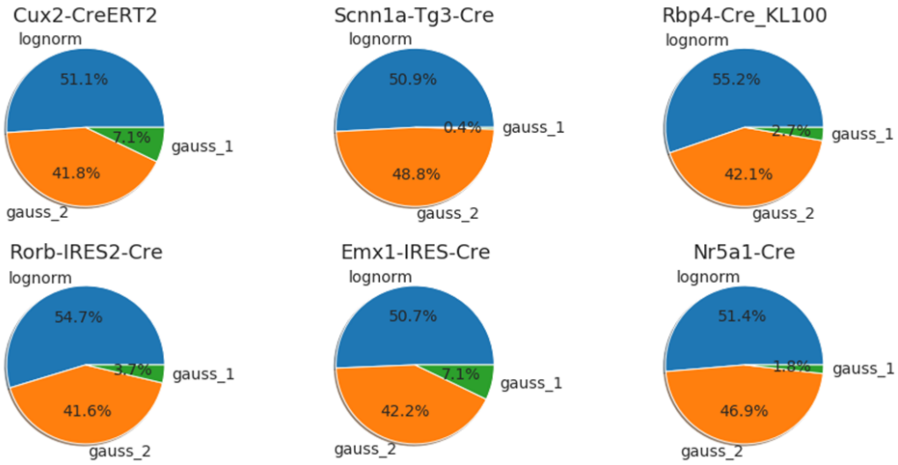


Figure 3: Dependence of noise distribution on cell types. We found relatively consistent results across all Cre-lines tested. The selected Cre-lines selectively label excitatory cells in different areas and layers.

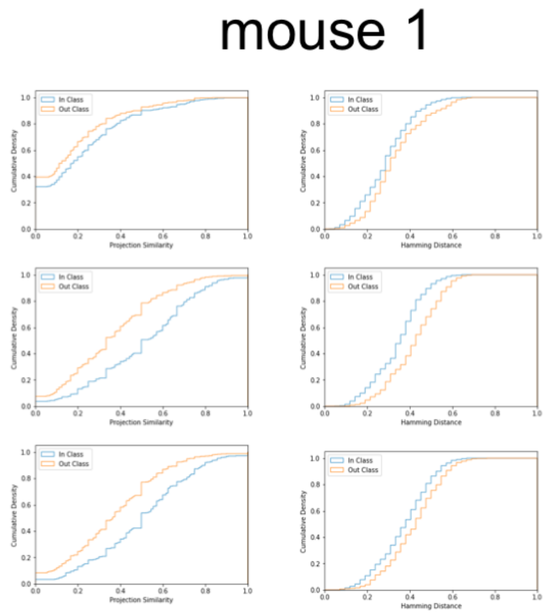


Figure 4: Cumulative distributions for similarity metrics using data aggregated across areas for mouse 1. Projection similarity (left column) and Hamming distance (right column) between clip variance subspace $V_k$ and noise subspace $N_{j,k,n}$ (line 1), between clip coding subspace $C_k$ and noise subspace $N_{j,k,n}$ (line 2) and between exemplar coding subspace $E_k$ and noise subspace $N_{j,k,n}$ (line 3). All the observed differences are statistically significant (KS test, $p < 0.05$).

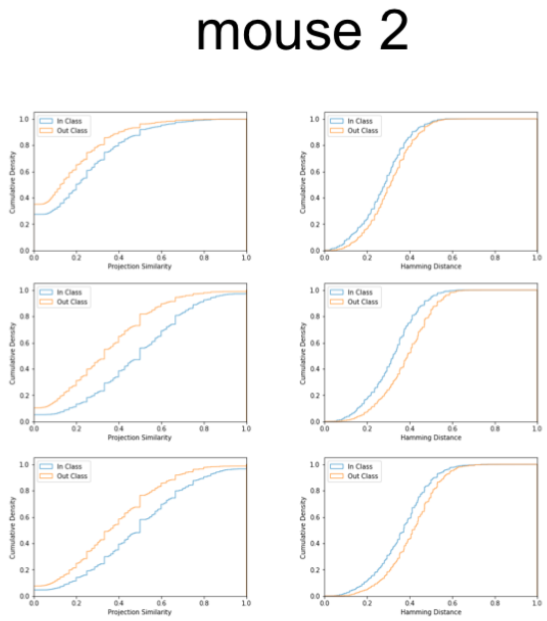


Figure 5: Cumulative distributions for similarity metrics using data aggregated across areas for mouse 2. Projection similarity (left column) and Hamming distance (right column) between clip variance subspace $V_k$ and noise subspace $N_{j,k,n}$ (line 1), between clip coding subspace $C_k$ and noise subspace $N_{j,k,n}$ (line 2) and between exemplar coding subspace $E_k$ and noise subspace $N_{j,k,n}$ (line 3). All the observed differences are statistically significant (KS test, $p < 0.05$).

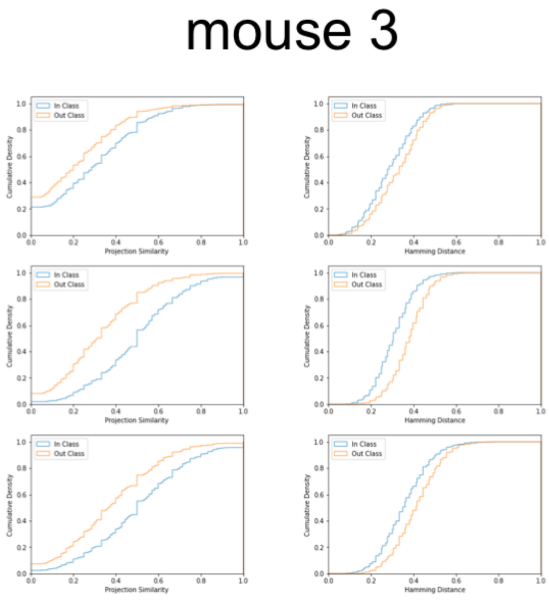


Figure 6: Cumulative distributions for similarity metrics using data aggregated across areas for mouse 3. Projection similarity (left column) and Hamming distance (right column) between clip variance subspace $V_k$ and noise subspace $N_{j,k,n}$ (line 1), between clip coding subspace $C_k$ and noise subspace $N_{j,k,n}$ (line 2) and between exemplar coding subspace $E_k$ and noise subspace $N_{j,k,n}$ (line 3). All the observed differences are statistically significant (KS test, $p < 0.05$).
